# Unraveling the diversity and functional potential of cyanosphere microbiomes assembled from terrestrial cyanobacteria

**DOI:** 10.1101/2025.06.10.658773

**Authors:** Brianne Palmer, Estelle M Couradeau, Jeffrey R. Johansen, Tania Kurbessoian, Jose Ortega Carranza, Jason E Stajich, Ryan Ward, Nicole Pietrasiak

## Abstract

The cyanosphere consists of heterotrophic microorganisms residing within the exopolysaccharide sheath of cyanobacteria, acting as a crucial interface between the cyanobacteria and their surrounding environment. Understanding the interactions between cyanobacteria and their cyanospheres is essential for predicting the success of terrestrial cyanobacteria in providing ecosystem services in nutrient-poor environments. However, knowledge of the microbial diversity within the cyanosphere remains limited.

In this study, we employed metagenomic sequencing to reconstruct 410 metagenome-assembled genomes (MAGs) from cyanosphere-associated microbes linked to 56 unialgal terrestrial cyanobacteria cultures, representing 12 distinct cyanobacteria orders. Our findings revealed that the composition of cyanosphere microbial communities was unique to each cyanobacterial host and was significantly shaped by environmental factors such as habitat, precipitation, and temperature from which the cultures were originally obtained. Notably, three microbial genera, *Brevundimonas*, *Devosia*, and *Sphingopyxis*, were present in over 30% of the cyanospheres, forming a core cyanosphere microbiome. Functional gene analysis showed a distinction between the cyanobacteria and their associated cyanospheres, with dissimilatory nitrate reduction being the dominant pathway in the cyanosphere, while nitrogen fixation was more common in the cyanobacteria. Three cyanospheres also contained nitrogen fixation genes of which two hosts were nitrogen fixation capable themselves. The cyanosphere harbored genes for polysaccharide lyases, indicating a possible link to the exopolysaccharides produced by the cyanobacteria. Given the observed variability in microbial community composition and function across different cyanobacterial hosts, future ecological assessments and restoration efforts involving cyanobacteria should not only focus on the cyanobacteria themselves but also consider their associated microbial communities.

**Importance:** Our study identifies members of a highly understudied, and potentially under-valued, microbial community -- the cyanosphere. We used a diversity of terrestrial cyanobacteria to understand how the cyanosphere composition and predicted functions were influenced by the host cyanobacterium and environmental factors using metagenomics. This is a new approach to study the cyanosphere and provides insights into the diversity of terrestrial microbial communities. Importantly, our results underscore the need to consider microbial consortia when assessing the ecological potential of cyanobacteria in terrestrial restoration.

## Introduction

Cyanobacteria are prevalent and perform various ecosystem functions in almost any environment where sunlight is at least temporarily available. This includes terrestrial environments which are highly complex and are characterized by often very contrasting abiotic and biotic conditions such as found in soil, hydroterrestrial, and subaerial habitats (1, 2). In soils, cyanobacteria contribute to carbon sequestration via organic and inorganic pathways (3), and enrich soil fertility by fixing atmospheric nitrogen and reducing soil erosion (4, 5). Cyanobacteria are also foundational components of biological soil crusts which cover approximately 30 % of drylands globally (6). In hydroterrestrial environments, such as periodically dry vernal pools, algal and cyanobacterial biofilms often form rapidly after rehydration and are important for maintaining local food webs as microscopic primary producers (7, 8). In subaerial habitats, which include exposed surfaces like rocks and tree bark, cyanobacteria are pioneers in colonizing these environments, contributing to the formation of epilithic and epiphytic biofilms that support other microbial and plant life (9–11). As early colonizers on these substrates, the cyanobacteria can enrich the substrate with organic carbon through photosynthesis and can improve nutrient cycling in the micro-environment by trapping dust and soil particles and fixing atmospheric nitrogen (10, 12).

A distinctive feature of many cyanobacteria is the presence of the cyanosphere (13). This is an ecological niche akin to the rhizosphere of plants. The cyanosphere is a microenvironment surrounding cyanobacterial cells, especially in filamentous forms, and is inhabited by heterotrophic microorganisms that closely associate with their cyanobacterial hosts (13). These heterotrophic microorganisms may benefit from the organic compounds produced by the cyanobacteria through photosynthesis, and in return, they can assist the cyanobacteria by providing essential nutrients (14). Mutualistic interactions are common in the rhizosphere and may thus also occur within the cyanosphere (15, 16).

Current research on the terrestrial cyanosphere is limited. Previous work on the cyanosphere has focused on *Microcoleus vaginatus*, a filamentous cyanobacteria known to be an early colonizer of biological soil crusts without the ability to fix nitrogen. In one of the first cyanosphere studies, Couradeau and her colleagues found that most of the inhabitants of the *M. vaginatus* cyanosphere were copiotrophs, and some were diazotrophs and contained more nitrogen-fixation genes in comparison to the bulk soil microbiome. This highlights the potential role of nutrient exchange of carbon and nitrogen between the cyanobacteria and the diazotrophs in the cyanosphere. A subsequent study found evidence of a carbon-nitrogen exchange between *M. vaginatus* and the cyanosphere (14). The success of *M. vaginatus* as an early colonizer and important species for cyanobacterial-mediated dryland soil restoration may be because of the heterotrophs within the cyanosphere such as *Arthrobacter, Massillia,* and *Bacillus* which, when added to the cyanobacterial growth medium, increased cyanobacterial growth (17). Nelson and his colleagues found evidence of signaling between *M. vaginatus* and its cyanosphere, further emphasizing the interconnectedness between the cyanobacteria and their associated heterotrophs (18).

Beyond *M. vaginatus,* a few studies have assessed the microbial community composition within the cyanosphere with most studies focused on either soil or freshwater habitats. In a direct comparison of the rhizosphere and the cyanosphere. Another study examined growth-promoting strains in both the rhizosphere and biocrust cyanospheres. They found 18 phyla common to both and identified growth-promoting isolates such as *Bosea* and *Pseudoarthrobacter* in the cyanosphere (16). Within aquatic environments, one study used metatranscriptomics to study the cyanospheres of both nitrogen-fixing and non-nitrogen-fixing aquatic cyanobacteria *Dolichospermum* and *Microcystis* showing that the bacterial associations were determined by the cyanobacterial host with the most abundant phyla being Bacteroidetes, Gemmatimonadetes, and Proteobacteria. They also identified functional redundancy within the cyanosphere (19). However, the diversity and ecology of hydroterrestrial and subaerial habitats are virtually unexplored.

Based on the limited research of terrestrial cyanospheres, we know that the identity of the microbes within the cyanosphere can be important for the success of the cyanobacteria (17), and the functions of the cyanosphere microbes can provide essential nutrients to the cyanobacteria (13, 14, 18). However, the breadth of terrestrial cyanobacterial species is large, and current research is limited to *M. vaginatus.* Therefore, there is a need to understand the identity and function of the cyanosphere microbes within the cyanosphere from a diverse set of terrestrial cyanobacteria.

The use of next-generation sequencing techniques allows for the identification of the microbial community within the cyanosphere. Metagenomics, in particular, provides information about the identity of the microbes and the potential functions they may be performing. Here, we used metagenomic sequencing on 56 cultured terrestrial cyanobacteria and their cyanosphere microbiomes. These cyanobacteria are classified into 12 of the currently recognized 19 orders in cyanobacteria and represent broad phylogenetic diversity. We sought to understand how the taxonomy of the host cyanobacteria and the environment affects the heterotrophic community of the cyanosphere.

## Methods

### Cyanobacteria cultures

Metagenomes were obtained from unialgal polycultures containing cyanobacteria and associated heterotrophic cyanosphere microbiomes. Polycultures were established via traditional dilution plating of environmental samples obtained from various collection trips led by Johansen, Pietrasiak, and collaborators. Collection years and detailed information can be found in Ward et al. (2021). Polycultures were isolated between 1965 and 2015 and represent reference cultures of recently well-studied taxonomic and phylogenetic benchmarks in cyanobacteria taxonomy and systematics. The dominant cyanobacterial members in these cultures are phylogenetically diverse, spanning 12 orders of cyanobacteria. Cultures have been maintained in two culture collections at John Carroll University (JCU) and University of Nevada - Las Vegas (UNLV) on solid or liquid Z8 media, an oligotrophic freshwater algal medium (20). Although dates of isolation span 50 years, close to identical culture conditions (use of Z8 medium, 16/8 hr light/dark photoperiod, low light intensity, grown in climate-controlled chambers) have been maintained at JCU since 1992 and have been reproduced at UNLV. At the time of DNA extraction, the cultures’ microbial biomass was dominated by cyanobacteria.

### Metagenomic Sequencing

Polyculture biomass was grown in liquid Z8 media, harvested, flash-frozen, and stored before DNA extraction using the Qiagen DNeasy PowerLyzer kit with bead-beating, following detailed protocols available on protocols.io (dx.doi.org/10.17504/protocols.io.brg4m3yw) (2). Extracted DNA was stored at -20°C prior to shipment to the Joint Genome Institute (JGI). JGI performed the library preparation and next gen Illumina sequencing steps as follows. DNA library preparation was performed using the KAPA Biosystems high-throughput library preparation kit p/n KK8235 on a PerkinElmer Sciclone next-generation sequencing (NGS) robotic liquid handling system. Then, 200 ng of sample DNA was sheared to 300 bp using a Covaris LE220 focused ultrasonicator. The sheared DNA fragments were size selected by double solid-phase reversible immobilization (SPRI). The selected fragments were end repaired, A-tailed, and ligated with Illumina-compatible sequencing adaptors from IDT containing a unique molecular index barcode for each sample library. The prepared libraries were quantified with a KAPA Biosystems quantitative PCR (qPCR) kit on a Roche LightCycler 480 real-time PCR instrument. Genomic libraries were sequenced with a NovaSeq instrument (Illumina, San Diego, CA) using NovaSeq XP V1 reagent kits and an S4 flow cell following a 2 x 150-bp indexed run recipe. Demultiplexed reads were processed with BBDuk v38.87 (17) to remove contaminants, trim adapter sequence and “G” homopolymers ≥5 in size at the ends, quality trim reads, and remove reads with ≥4 “N” bases, with an average quality score of >3, or with a length of <51 bp. Raw sequence products were submitted to NCBI SRA archive. A list of BioProjects is provided in Ward et al. (2) with additional descriptions in Table S1.

### Microbial community analyses

Sequence Read Archive (SRA) reads were first uploaded into the KBase platform (21). The quality of these reads was assessed using the Fast-QC tool (22), and those that did not meet the quality threshold (Q score > 30) were refined using Trimmomatic (23). The reads for each Cyanobacteria were assembled using metaSPAdes (24). The assembled contigs were then segregated into Metagenome-Assembled-Genomes (MAGs) via MaxBin2 (25). Taxonomic classification was conducted using GTDB-tk (26). A comprehensive taxonomy table, incorporating DRAM annotation, was generated to establish a connection between taxonomy and function. MAGs that did not meet the minimum requirements for medium-quality MAGs (> 50% completion and < 10% contamination) (27) were removed. The KBase narrative is publicly available at https://narrative.kbase.us/narrative/202318.

All subsequent analyses were performed in R (version 4.3.1). The *amp_alpha_diversity* function within the R package *ampvis2* was used to calculate alpha diversity metrics including the total number of MAGs and Shannon diversity (28). Differences in MAG richness and Shannon diversity between categorical variables were calculated using ANOVAs with Tukey HSD post-hoc tests.

To determine if there is a core cyanosphere, we calculated the number of cyanospheres in which heterotrophic genus occurred. Genera found in > 30% of the cyanospheres were classified as part of the core cyanosphere. The package *microeco* was also used to create a heatmap of the most abundant genera.

Bray-Curtis distances were used to calculate the beta diversity using the *microeco* package. To assess differences in the microbial community composition across categorical variables, a PERMANOVA analysis was performed using the *cal_manova* function. Beta diversity was visualized through Principal Coordinates Analysis (PCoA) plots generated in *microeco* R package (29).

Climate variables were obtained from BIOCLIM (30). When latitudinal and longitudinal data were available, we identified the mean annual temperature and the annual precipitation at the collection location of each cyanobacterium in BIOCLIM. Due to the covariance between environmental variables, only mean annual temperature and precipitation were used. The impact of continuous variables such as temperature, precipitation, and altitude on microbial community composition was evaluated using the *cal_ordination_envfit* function within *microeco*. This relationship was further illustrated with a distance-based Redundancy Analysis (dbRDA) plot using the *microeco* R package (29). The functional abundances, as assigned via DRAM, were visualized using heat maps created with the *pheatmap* R package (31) for the cyanosphere community and the host Cyanobacteria. The code for this project is publicly available on GitHub at https://github.com/briannepalmer/Cyanosphere.

## Results

### Spatial representation

Cyanobacterial hosts represented 14 locations spanning 4 continents (including the Pacific Islands) (Figure 1). Soil Cyanobacteria were collected from 5 types of parent material (dolomite, granitic, gypsiferous, gneiss, and limestone) (Table S1). The altitude of the host Cyanobacteria ranged from 11 to 2043 meters. The annual precipitation was between 3 mm and 3229 mm. The annual mean temperature ranged from 5.2 °C to 23.6 °C. In total, 410 cyanosphere-associated microbial MAGs were identified, 214 from soil cyanobacteria, 159 from subaerial cyanobacteria, 23 from freshwater cyanobacteria, and 13 from hydroterrestrial cyanobacteria.

**Figure 1:**
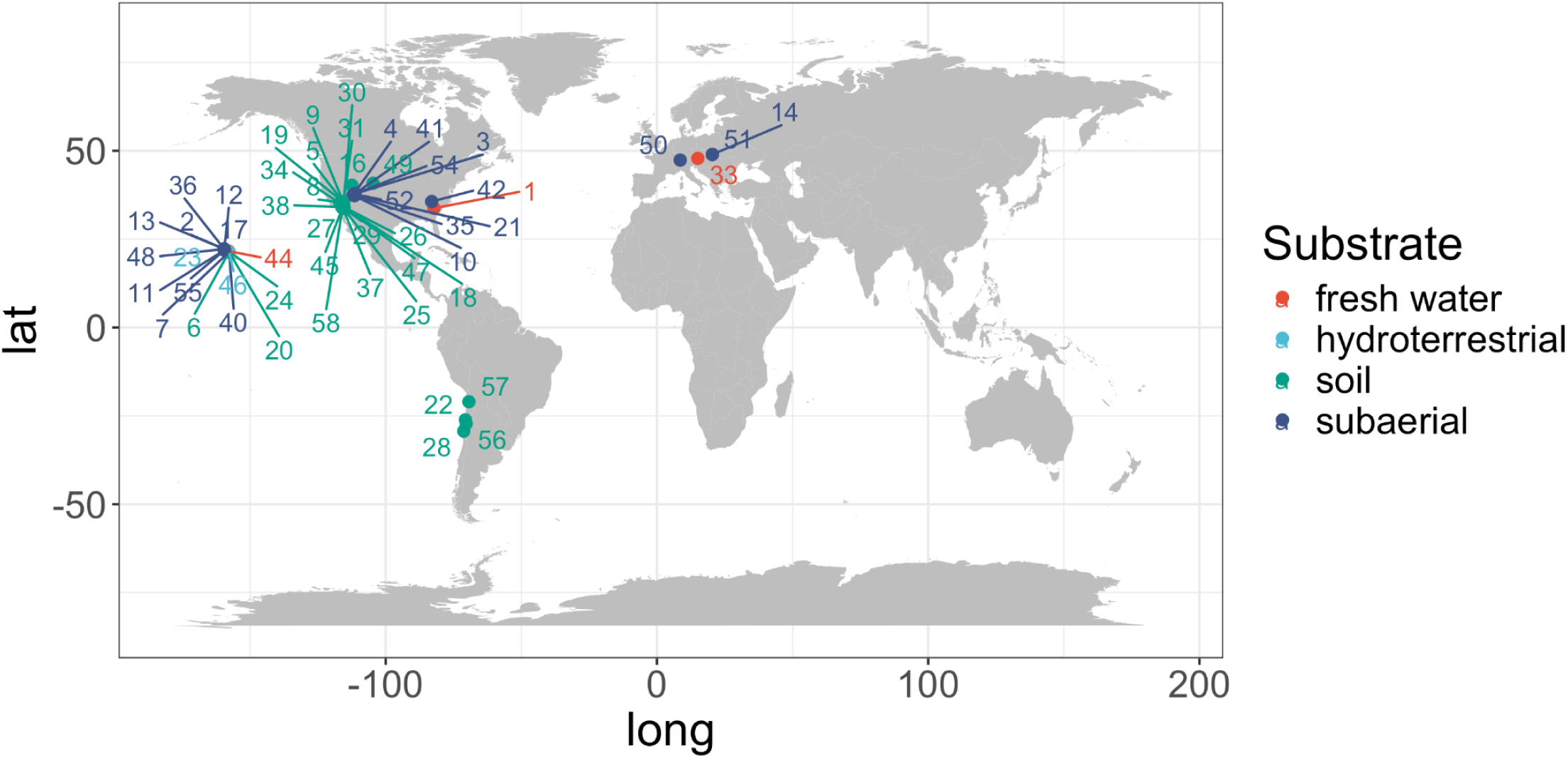
Map of the locations where the cyanobacteria were collected. Five cyanobacteria (*Drouetiella lurida, Oculatella coburnii, Pleurocapsa ergoicii, Roholtiella mojaviensis* and *Timaviella radians*) did not have coordinate data and are not included on the map. The points are colored by the habitat where the cyanobacteria were collected, and the numbers are assigned to each cyanobacterium found in Table S1.

### Core and Most Abundant Genera

Proteobacteria (72%), Bacteroidota (11%), and Actinobacteria (6%) were the most abundant phyla in the cyanospheres (Figure 2A). Within Proteobacteria, Rhizobiales was the most abundant order, occurring in 79% of the cyanospheres (Figure 2B). Only three genera were found in > 30% of the samples: *Brevundimonas* (40%)*, Devosia* (36%), and *Sphingopyxis* (30%) (Figure 3). These genera could represent a core terrestrial cyanobacterial cyanosphere. There were two families in > 50% of the cyanospheres: Sphingomonadaceae and Caulobacteraceae. No genera were found in every cyanobacterial order. However, two genera – *Allorhizobium* and *Sphingopyxis* – were found in all four habitat types. Additionally, *Brevundimonas, Devoisa,* and *Sphingomonas* were found in the three terrestrial habitats (soil, subaerial, hydro-terrestrial).

**Figure 2:**
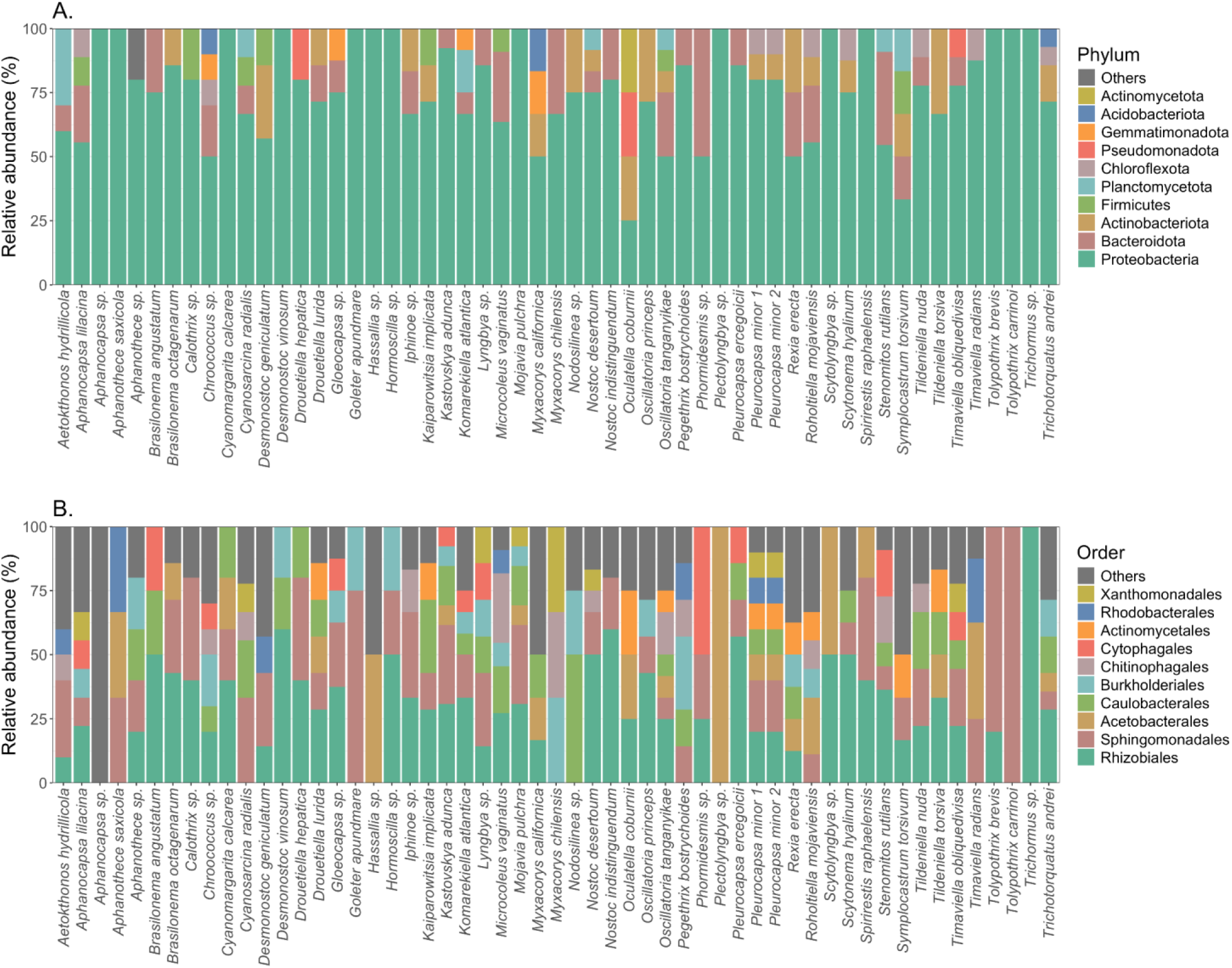
Bar plots depicting the relative abundance of the most abundant phyla (A) and the Proteobacteria orders (B) for the MAGs in the cyanosphere.

**Figure 3:**
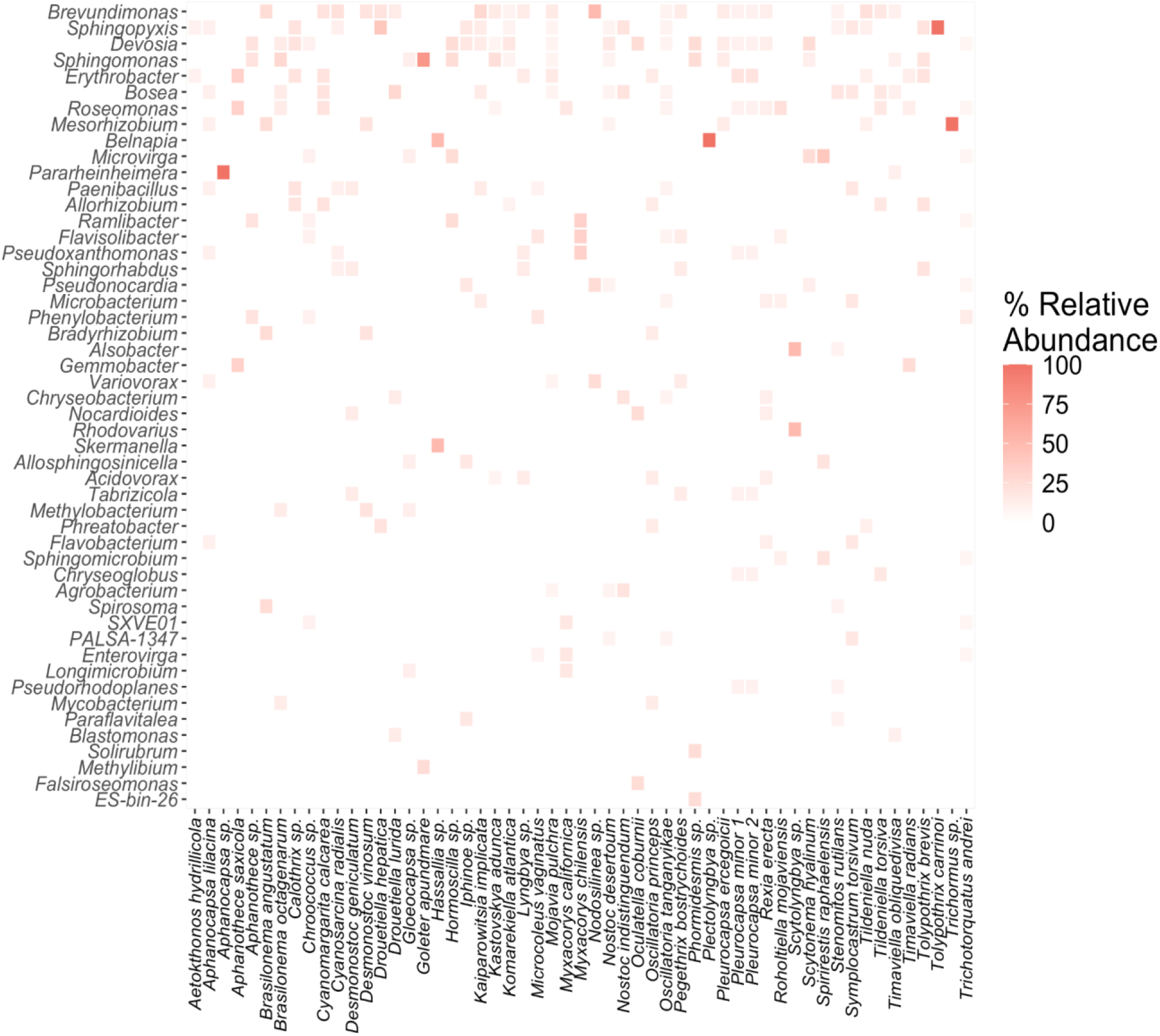
Heatmap showing the relative abundance of the 50 most abundant heterotrophic genera found within each cyanobacterial host. Dark red indicates higher relative abundance while light red and white indicates low or no abundance in the given cyanosphere. The heterotrophic genera on the y-axis are ordered by the total relative abundance across all cyanospheres (x-axis).

### Alpha Diversity Metrics were Consistent Across Cyanobacterial Hosts

MAG richness and Shannon diversity did not vary among cyanobacterial orders (ANOVA, P = 0.18, P = 0.06) or among habitats (ANOVA, P = 0.61, P = 0.54) (Figure 4). *Chroococcus* sp., *Trichotorquatus andrei,* and *Mojavia pulchra* each had the most heterotrophic MAGs (n = 14) followed by *Kastovskya adunca, Oscillatoria tanganyikae,* and *Microcoleus vaginatus* (n = 13). *Pelatocladus maninholoensis* did not contain any non-cyanobacterial MAGs. *Plectolyngbya* sp., *Trichormus* sp., *Scytonematopsis contorta,* and *Aphanocapsa* sp. only contained 1 non-cyanobacterial MAG. There was an average of 8 MAGs per cyanobacteria.

**Figure 4:**
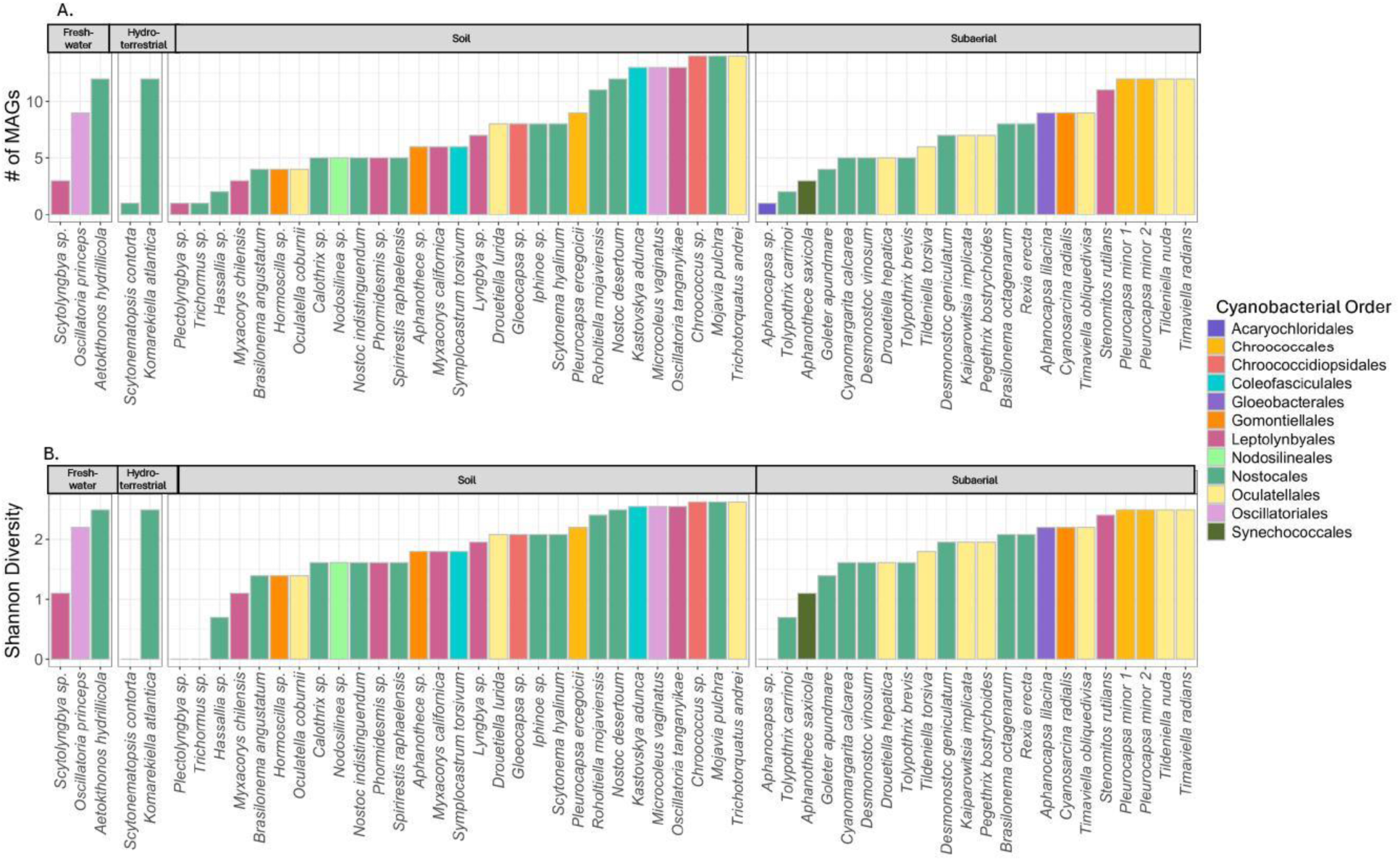
Bar charts depicting the total number of MAGs (A) and the Shannon diversity of the MAGs (B) for each Cyanosphere, grouped by habitat. The bars are colored by the order of the host Cyanobacteria.

### Microbial Community Composition Varies Between Habitats, Rather than Between Cyanobacterial Hosts

Based on the PcOA plot and PERMANOVA (P = 0.08), there are no differences in the heterotrophic community composition among cyanobacterial host orders. The PCoA axis explain 12.7% and 9.6% of the variation in the community composition and there is little clustering between heterotrophic communities with the same cyanobacterial host order (Figure 5A). However, the microbial community composition did vary by host genera (PERMANOVA, P = 0.02) highlighting that each cyanosphere contains a unique microbial community. There was also a significant effect of habitat on the community composition (PERMANOVA, P = 0.001). The mean annual temperature (P < 0.001), annual precipitation (P < 0.001), and altitude (P 0.015) where the cyanobacteria were collected also influenced the community composition (Figure 5B). When the environmental factors and habitat type are included in an RDA plot, there is distinct grouping between the soil and subaerial samples and the axes explain 46.8% and 31.6% of the variation in the microbial community composition. The soil communities are grouped with the annual mean temperature vector, while the subaerial samples group with annual precipitation.

**Figure 5:**
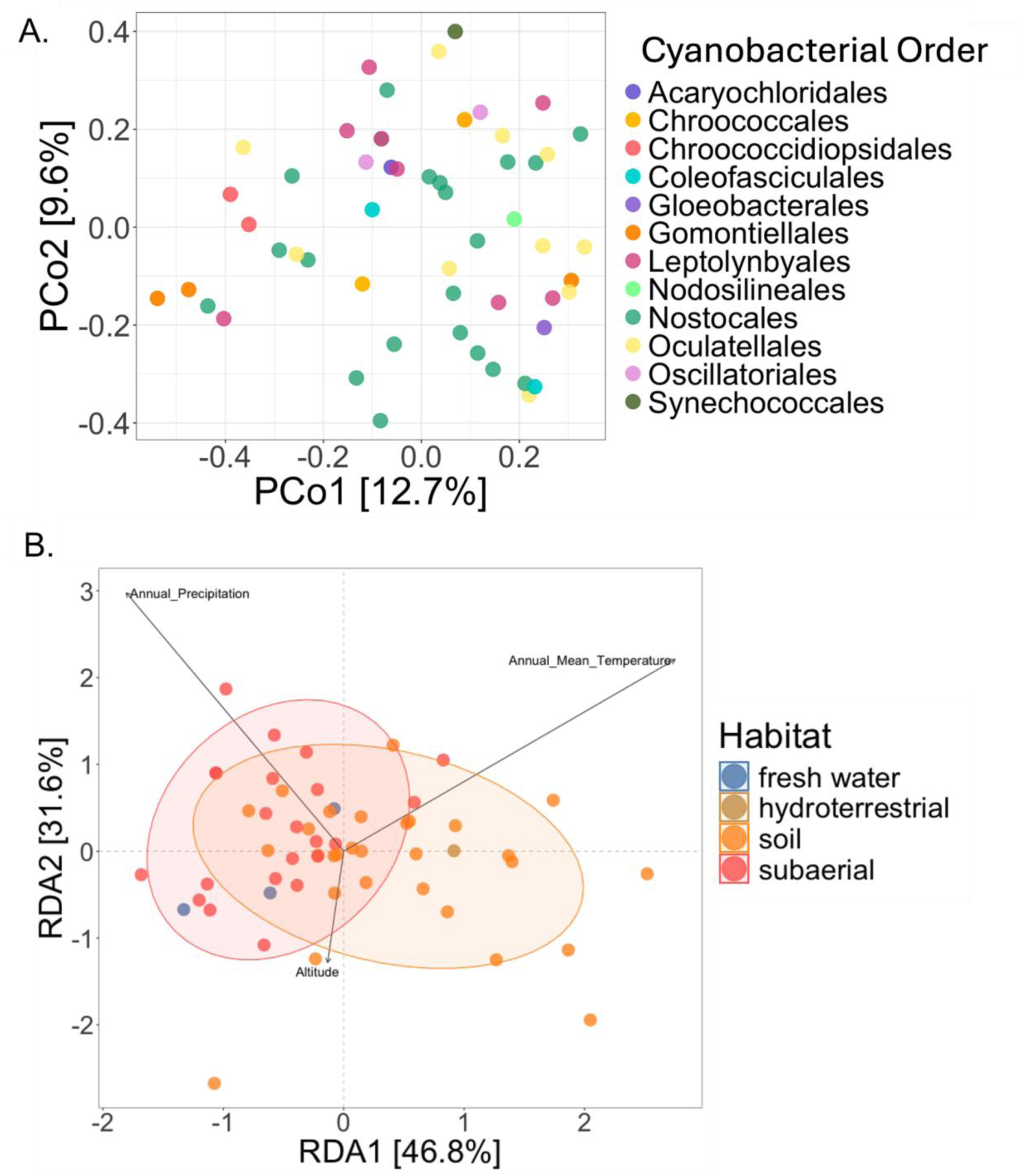
Ordination plots showing the community composition based on the MAG genera in the cyanospheres. A) a PCoA depicting the MAG community composition colored by the order of the cyanobacterial host. B) a RDA plot colored by habitat indicating the grouping of the cyanosphere community composition. The ellipses show groupings of the cyanospheres with more than 3 representatives. The arrow vectors (Annual Precipitation, Annual Mean Temperature, and Altitude) show environmental variables that were significantly (P<0.05) correlated with the cyanosphere community composition.

### Cyanosphere and cyanobacterial hosts differ in key functions

Only three cyanospheres contained cyanosphere members with nitrogen-fixation genes: *Hassallia* sp., *Brasilonema angustatum,* and *Oscillatoria princeps* (Figure 6). These putative diazotrophs included *Bradyrhizobium* in *Brasilonema angustatum* and *Oscillatoria princeps*, as well as *Skermanella* in *Hassallia* sp.

**Figure 6:**
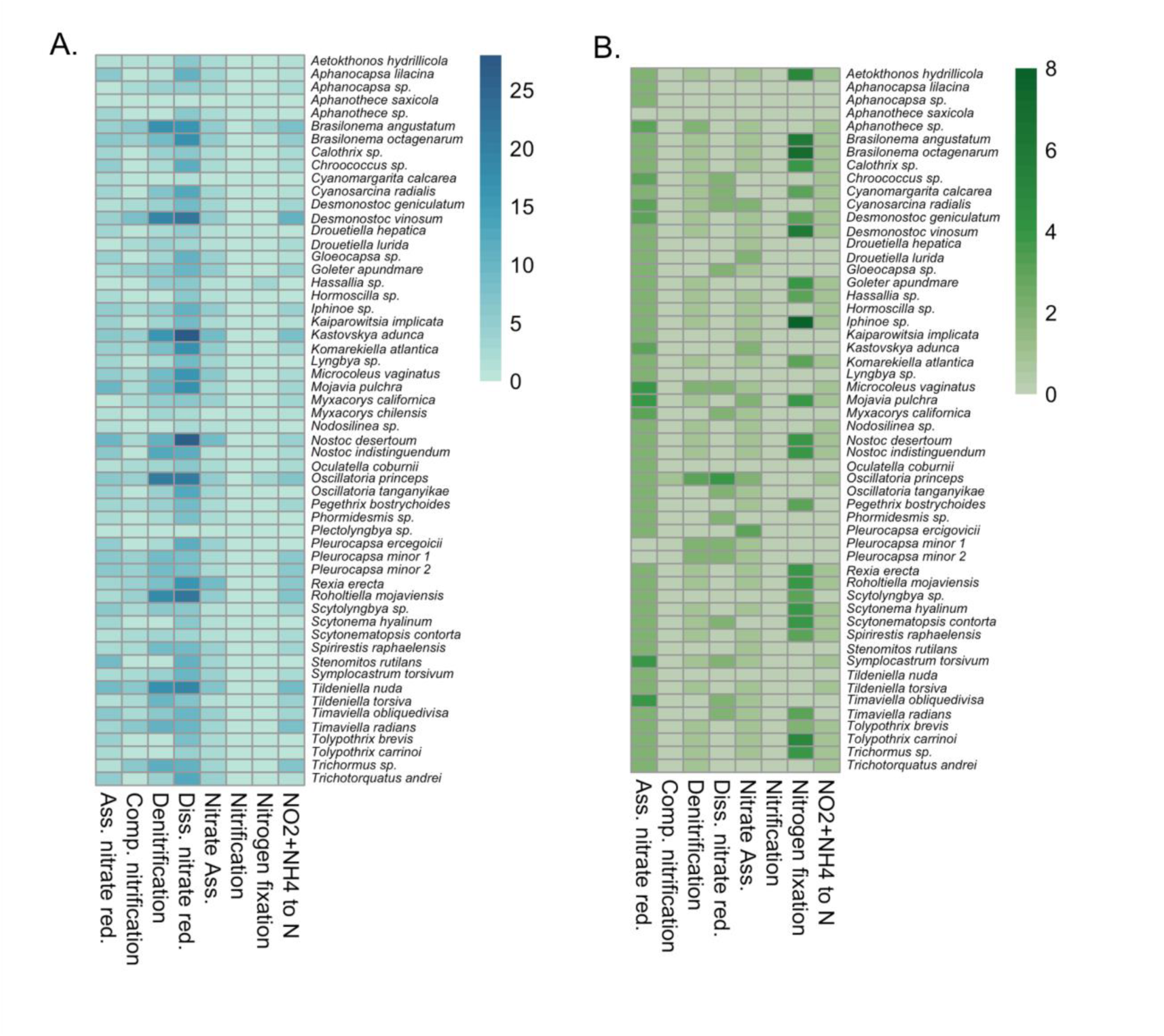
Heatmaps depicting the abundance of genes relating to nitrogen pathways for the cyanosphere (A) and cyanobacteria (B). The x-axis labels describe the nitrogen pathways including assimilatory nitrate reduction, complete nitrification, denitrification, dissimilatory nitrate reduction, nitrate assimilation, nitrification, nitrogen fixation, and nitrite + ammonia => nitrogen.

Dissimilatory nitrate reduction emerged as the most prevalent nitrogen pathway within the cyanosphere (Figure 6), although it was notably absent in the cyanospheres of *Aphanothece saxicola* and *Plectolyngbya* sp. The cyanospheres of *Kastovskya adunca* and *Nostoc desertorum* had the highest number of genes coding for dissimilatory nitrate reduction (DNRA), while denitrification was most prominent in the cyanosphere of *Oscillatoria princeps*. Genes for assimilatory nitrate reduction were most abundant in the cyanospheres of *Nostoc desertorum* and *Mojavia pulchra*, whereas *Desmonostoc vinosum* exhibited the most complete nitrification pathways. Nitrate assimilation was most pronounced in cyanosphere microbes of *Kastovskya adunca*, *Nostoc desertorum*, and *Rexia erecta*. Notably, none of the cyanospheres contained pathways for nitrification.

This pattern contrasted with the nitrogen pathways observed in the cyanobacteria themselves. *Myxaxorys chilensis*, *Plectolyngbya* sp., and *Trichocoleus desertorum* lacked high-quality cyanobacterial MAGs, while *Timaviella obliquedivisa* had two cyanobacterial MAGs. *Iphinoe* sp. contained the highest number of genes for nitrogen fixation, followed by *Brasilonema octagenarum* and *Brasilonema angustatum* (Figure 6). In total, 25 cyanobacterial species possessed nitrogen fixation genes, most corresponding with the order Nostocales. Both *Hassallia* sp. and *Brasilonema angustatum*, whose cyanosphere microbes had nitrogen fixation genes present contained cyanobacterial nitrogen fixation genes corresponding to their ability to produce heterocytes and thus taxonomic affiliation to the order Nostocales taxa, while *Oscillatoria princeps*, order Oscillatoriales, did not contain nitrogen fixation genes. Interestingly, cyanobacteria taxonomically not affiliating with the order Nostocales also contained nitrogen fixation encoding genes, such as, *Pegethrix bostrychoides*. As with the cyanospheres, no genes involved in nitrification were detected in the cyanobacteria.

Carbohydrate-Active enZYme (CAZy) pathways were also prevalent in the cyanospheres, particularly those involving glycosyl transferases and glycoside hydrolases (Figure 7). The cyanospheres of *Chroococcus* sp. and *Roholtiella mojaviensis* contained the highest number of genes related to glycosyl transferases, while *Aphanocapsa* sp. had the fewest. For glycoside hydrolases, *Microcoleus vaginatus* and *Chroococcus* sp. cyanospheres had the most genes, whereas *Scytonematopsis contorta* had the fewest. Although polysaccharide lyases were less abundant, they were still detected in 48 cyanospheres, with genes for polysaccharide lyases present in 68 genera (Figure 7). The *Nostoc desertorum* cyanosphere contained the most polysaccharide lyase genes, while *Scytolyngbya* sp. had only one.

**Figure 7:**
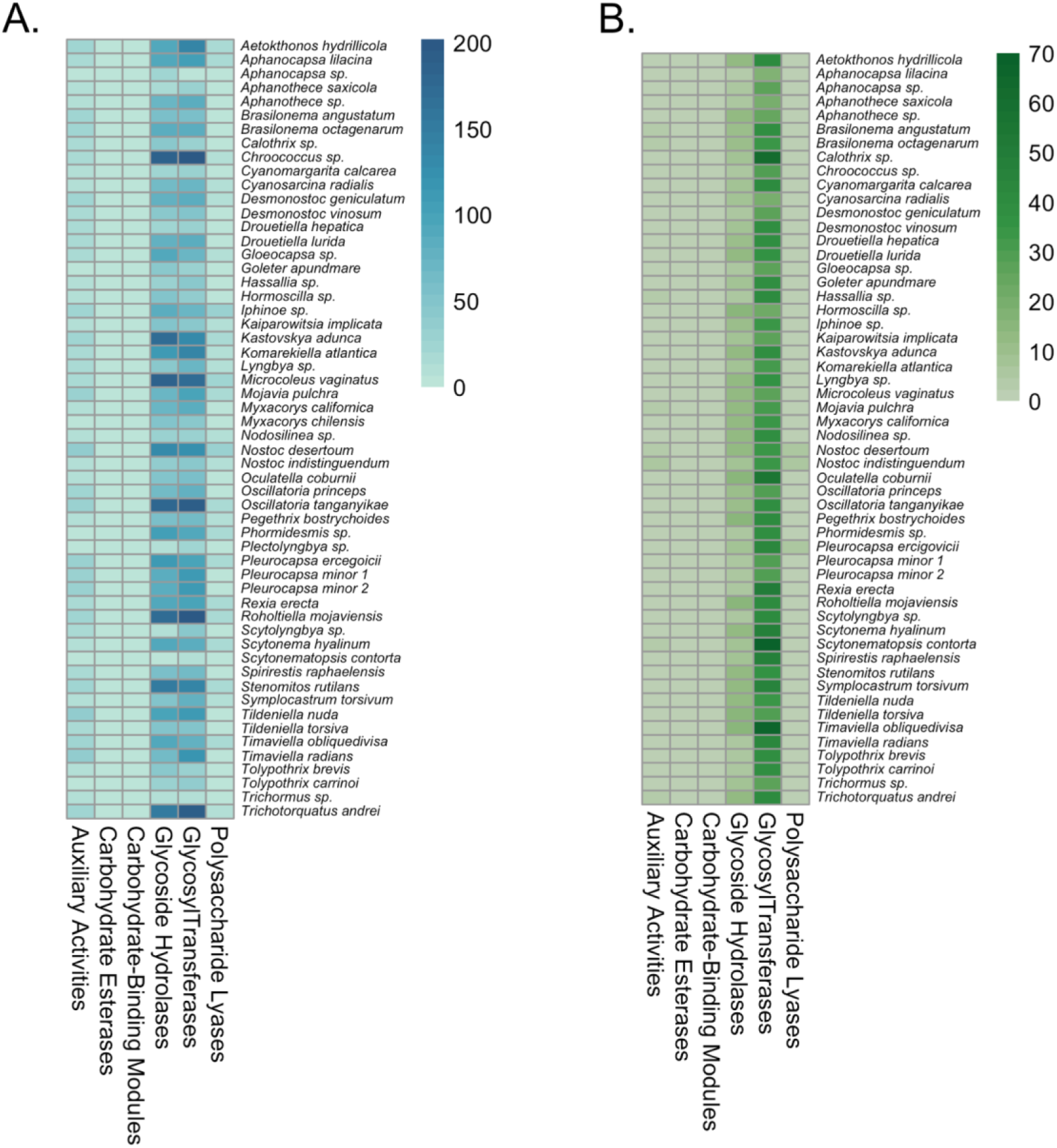
Heatmaps depicting the abundance of genes relating to CAZy pathways found in the cyanosphere (A) and the cyanobacteria (B). The genes present in the cyanosphere differ notably from those found in the cyanobacterial host. In the cyanobacteria, *Scytonematopsis contorta* had a high abundance of glycosyl transferase genes, while *Timaviella obliquedivisa* possessed the most glycoside hydrolase-related genes (Figure 7). Polysaccharide lyases, however, were relatively scarce in cyanobacteria, with only 24 species containing these genes. Of those, only *Nostoc indistinguendum*, *Nostoc desertorum*, *Pleurocapsa ercegoicii*, *Roholtiella mojaviensis*, and *Scytolyngbya* sp. had more than one gene associated with polysaccharide lyases.

Interestingly, genes for carbohydrate-binding molecules and carbohydrate esterases were absent from all cyanospheres. Among the polysaccharide lyases identified, five alginate lyases were found—PL5, PL6, PL15, PL17—all capable of degrading alginate. Thirty cyanospheres, mostly from *Nostocales* (n=14), contained alginate lyases, while *Synechococcales* cyanospheres lacked these enzymes (Figure 8). The *Iphinoe* sp. cyanosphere had the highest number of alginate lyases, followed by *Microcoleus vaginatus*.

**Figure 8:**
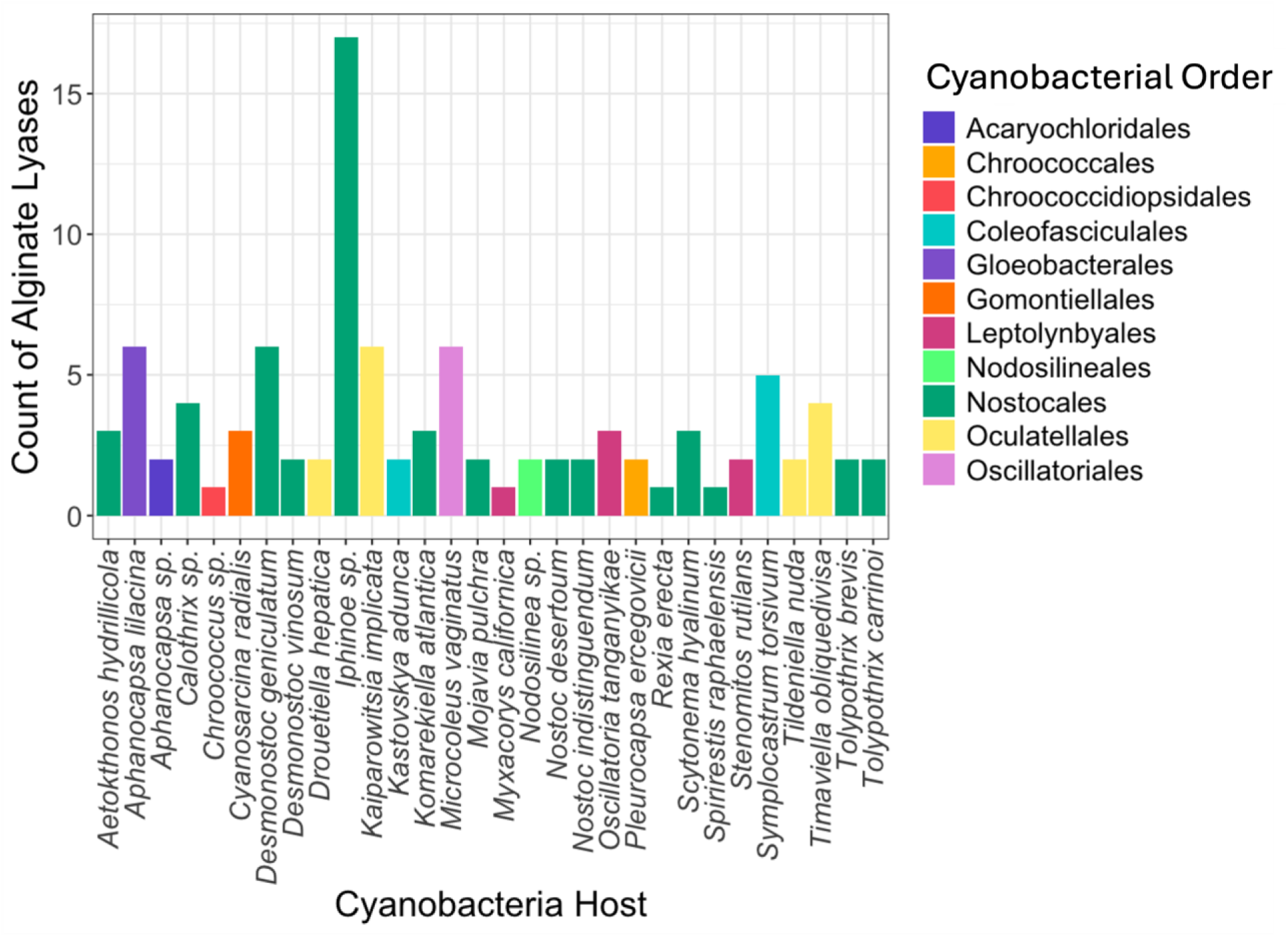
The total number of alginate lyases found in the cyanosphere of each cyanobacteria host. The colors represent the order of the cyanobacterial host.

## Discussion

Using metagenomic sequencing, we identified non-cyanobacterial members of the cyanosphere from various habitats and cyanobacterial orders. This study is the first, to our knowledge, to identify the cyanosphere for a broad scope of terrestrial cyanobacteria and has implications for understanding interspecific interactions and the growth conditions necessary for cyanobacteria.

### Trends within cyanospheres

Based on 410 non-cyanobacteria heterotrophs from 56 cyanobacterial hosts, we identified patterns between the cyanospheres. The most common genera, found in over 30% of the cyanospheres were *Brevundimonas, Devosia,* and *Sphingopyxis.* These genera might indicate a “core microbiome” as prior studies have used abundance in 30% of samples as an arbitrary cutoff for calculating core communities (32, 33). *Brevundimonas* is a diverse genus and species have been found in both soil and aquatic environments (34–36). In the present study, *Brevundimonas* were found in soil, subaerial, and hydroterrestial environments and accounted for 6.8% of the MAGs in the cyanosphere. *Brevundimonas* species can live as free-living species in a variety of environments including as a diazotrophic plant-root colonizer (37), the aerobic and anaerobic conditions of activated sludge (38), alkaline soils (39), and thermal baths (40). The ability to thrive in such variable environmental conditions may be beneficial in the cyanosphere, as the cyanobacteria were collected from diverse environments.

*Devosia* is a common genus in soil and aquatic sediment (41–45). It was also relatively common in the cyanospheres, found in 36% of the cyanospheres and representing 5.6% of the total MAGs. At least one species of *Devosia* can produce nitrogen-fixing root-nodule symbiosis with plants (46), indicating nitrogen fixation capability, which may also occur in the cyanosphere when nitrogen fixation is needed. However, there were no nitrogen-fixation genes detected in the cyanosphere *Devosia* in the present study. Two species of *Devosia* were previously isolated from the phycosphere of marine red alga (47). Previous work highlighted the genomic plasticity of *Devosia*, which allows this genus to utilize a wide variety of substrates (48). This ability to adapt to different environments may contribute to the prevalence of *Devosia* in the cyanosphere.

In the cyanosphere, *Sphingopyxis* was found in 30% of the host cyanobacteria and represented 4.4% of the total cyanosphere MAGs. Like *Devosia, Sphingopyxis* is a widespread genus due to its ability to thrive in diverse and often stressful environments such as aquatic sediment (49), contaminated soil (50, 51), and activated sludge (52). Likely contributing to its survival in stressful environments is the ability of several species to degrade environmental contaminants and toxins and produce secondary metabolites (53).

The order, Rhizobiales was present in 79% of the cyanospheres, indicating that this order is an important constituent. Rhizobiales contains several nitrogen-fixing genera, primarily associated with root nodules, like *Rhizobium.* This is a broad taxonomic group, and not all genera can fix nitrogen. Interestingly, several Rhizobiales known to fix nitrogen were found in the cyanosphere, despite living in the cyanosphere of a nitrogen-fixing cyanobacteria including: *Bosea, Mesorhizobium, Devosia, Bradyrhizobium, Methylobacterium, Allorhizobium, Microvirga, Ensifer, Neorhizobium, Agrobacterium, Shinella,* and *Pararhizobium* (54, 55). These genera are known root-nodule formers.

However, we did not detect *nif* genes in their MAGs and currently we do not know if these microbes have an established symbiotic relationship with the cyanobacteria for carbon and nitrogen exchange like in plant roots. Rhizobiales that do not fix nitrogen were also found in these cyanospheres. These may be pathogenic, similar to the Rhizobiales root pathogens found in the rhizosphere (56). In the present study, only three cyanosphere microbes had nitrogen-fixing genes. This indicates that perhaps in most cases, the cyanosphere selects for members without the ability to fix nitrogen, or the metagenomes do not fully represent the metabolic diversity of the cyanosphere.

The richness and diversity of the microbial community did not vary significantly between the cyanospheres while composition was influenced by the cyanobacterial host. A previous study with the rhizosphere showed that bacterial richness and diversity were greater in the bulk soil compared to the rhizosphere and that the richness and diversity of the rhizosphere rarely varied between rhizospheres (57). The authors state that the rhizosphere microbiota have similar traits regardless of the plant, thus resulting in similar richness and diversity measurements regardless of the phylogenetic placement of the bacteria. A similar phenomenon may occur in the cyanosphere, whereby bacteria filling particular niches colonize the cyanosphere regardless of cyanobacterial host phylogeny resulting in similar measurements of richness and alpha diversity between cyanospheres.

### Habitat influences cyanosphere microbiome

The community composition of the cyanosphere was influenced more by the habitat where the cyanobacteria were collected than the phylogeny of the cyanobacteria. This suggests that the microbes forming the cyanosphere are not determined by relationships between the cyanosphere and the phylogenetic affiliation of the host cyanobacteria at the broader taxonomic level.

The effect of habitat could be due to recruitment of the cyanosphere microbiome from the surrounding microbial community. This commonly occurs within the rhizosphere where the rhizosphere microbiome is a specific subset of the bulk soil microbiome (58). In the rhizosphere, root traits can explain the community composition (59). Thus, in the cyanosphere, the traits of the cyanobacteria could be the factors that determine which bacteria colonize the cyanosphere.

One of the limits of this dataset is the lack of even sampling across locations. This should be rectified in future studies. As such, there was a bias between habitat and sampling location. For example, 21 of the 29 soil cyanospheres were collected from North America (primarily Mojave and Sonoran Deserts, Southern California). Thus, it is difficult to parse out if the difference in the community composition between habitats is due to the effect of the particular habitat or due to the effect of sampling location, in this case, with a bias towards microbial communities that are present in California. The problem is not persistent within the other habitat types as the subaerial samples (22) have 3 from Europe, 10 from North America, and 9 from the Pacific Islands and the freshwater samples are from Europe, North America, and the Pacific Islands. Furthermore, there was not even sampling across habitats with 29 samples from the soil, 22 subaerial, 3 fresh water, and 2 hydroterrestrial (both from Hawaii). This should not discount the results presented in our exploratory study thus far but rather should be used as a stepping stone for subsequent research. Given the differences in the microbial community composition between habitats (or locations), there is a need for further research to determine the specific factors that influence the cyanosphere microbial community composition.

### Cyanospheres and cyanobacteria differ in predicted nitrogen and carbon exchange pathways

Unlike the rhizosphere, there is not a clear relationship of carbon for nitrogen exchange between the cyanosphere and the host cyanobacteria, perhaps because many cyanobacteria can fix nitrogen on their own, including 25 Cyanobacteria included in this study. However, three cyanospheres contained nitrogen-fixing genes within the cyanosphere: *Hassallia* sp., *Brasilonema angustatum,* and *Oscillatoria princeps. Brasilonema angustatum* and *Hassallia* sp., both of the order Nostocales, contain nitrogen-fixation genes in their genomes, are able to form heterocytes, and can fix atmospheric nitrogen (60, 61). Thus, these cyanobacteria would not necessarily need to interact with nitrogen-fixers in their cyanosphere. Thus, it is puzzling why we found diazotrophic *Bradyrhizobium* and *Skermanella* in the cyanosphere of these two diazotrophic cyanobacteria. *Bradyrhizobium* is a common genus in the rhizosphere and can form root nodules (62, 63), and both *Skermanella* and *Bradyrhizobium* were identified as key indicator species of potential diazotroph activity in terrestrial ecosystems (64). Perhaps under certain environmental conditions either the cyanobacteria or the cyanosphere bacteria (*Bradyrhizobium* and *Skermanella*) are not performing nitrogen fixation and gaining nitrogen through the symbiotic relationship. In comparison, *Oscillatoria princeps* did not have genes for nitrogen fixation, does not form heterocytes but recorded a *Bradyrhizobium* MAG in its cyanosphere, suggesting a possible mutualistic relationship with *Bradyrhizobium* in the cyanosphere.

Neither the Cyanobacteria nor the cyanospheres had genes for nitrification. The nitrogen metabolism genes in the Cyanobacteria emphasize their role in assimilatory nitrate reduction and nitrogen fixation; while the cyanosphere has more genes related to dissimilatory nitrogen reduction and denitrification. Dissimilatory nitrate reduction reduces nitrate to nitrite through denitrification then reduces nitrite to ammonium, conserving nitrogen as ammonium rather than dinitrogen gas. Assimilatory nitrate reduction reduces nitrate to nitrite and ammonium which can be incorporated into the organism. The prevalence of these two different pathways of nitrogen reduction indicates the different nitrogen priorities of the cyanosphere and host cyanobacteria.

Within the CAZy framework, the abundance of polysaccharide lyases in the cyanosphere is notable, as this may provide a mechanism for cyanobacterial EPS degradation or consumption (65). The EPS could provide a source of carbon for the cyanosphere. Polysaccharide lyases were found in 86% of the cyanospheres, highlighting the importance of this gene family within cyanospheres across cyanobacteria habitats. The key cyanosphere genera, *Sphingomonas* and *Brevundimonas,* are known for containing polysaccharide lyases. *Sphingomonas* strains can synthesize gellan lyase (66) and alginate lyases (67). *Sphingomonas* with polysaccharide lyase genes was found in 7 of the cyanospheres and was present in all terrestrial environments (1 subaerial, 1 hydroterrestrial, 5 soil). Perhaps its ability to degrade EPS contributes to its prevalence in all three terrestrial habitats. *Brevundimonas*, part of the “core” cyanosphere, also contained several polysaccharide lyases within 28% of the cyanospheres with the greatest abundance in *Kastovskya adunca, Mojavia pulchra,* and *Nodosilinea* sp. A particular group of polysaccharide lyases – alginate lyases– can degrade alginate, a polysaccharide found in algae and cyanobacteria (68, 69). The cyanosphere contained six types of alginate lyases (PL6 alginate lyase, PL5 alginate lyase, M-specific alginate lyase, PL15 oligo-alginate lyase, PL17 alginate lyase, PL14 alginate lyase).

Interestingly, polysaccharide lyases were also found in 42% of the cyanobacteria, with the most genes in *Nostoc indistinguendum, Nostoc desertorum,* and *Pleurocapsa ercegovicii. Nostoc*, in particular, is known for containing and utilizing polysaccharide lyases. Plant symbiotic *Nostoc* strains associated with feathermoss contained pectate lyases (70) which can be involved in plant pathogenesis and allow for symbiont-rhizobia infection (71, 72). The pectate lyases found in feathermoss-associated *Nostoc* may assist with the colonization by *Nostoc* of the feathermoss. In the cyanosphere, this process would need to be reversed with various heterotrophic bacteria colonizing *Nostoc*.

### Implications

The application of terrestrial cyanobacteria in ecosystem restoration is an emerging and rapidly expanding field (73–75). These microorganisms play a vital role in dryland ecosystem restoration due to their remarkable ability to withstand environmental stressors and the various ecological functions they support (76). Cyanobacteria contribute to critical processes such as stabilizing soils, enhancing soil fertility, promoting nutrient cycling, increasing water retention and infiltration, and sequestering carbon (74, 75, 77, 78). These functions are essential for restoring degraded environments, particularly in arid and semi-arid regions where soil erosion, nutrient depletion, and poor water retention are prevalent challenges.

The success of restoration efforts involving cyanobacteria can be closely tied to the cyanosphere (17). The composition of the cyanosphere may play a crucial role in determining the effectiveness of cyanobacteria in restoration activities. Therefore, it is essential 1) to investigate the relationships between cyanosphere microbes and their host more deeply, and 2) to consider the unique characteristics and requirements of the cyanosphere when selecting and applying cyanobacteria in ecosystem restoration projects to optimize their performance and ensure long-term ecological benefits.

## Data Availability

The sequence reads for the project are deposited under BioProjects PRJNA1200267, PRJNA1208396, PRJNA1208397, and PRJNA1231202 (Table S1).

## Acknowledgments

We are grateful to the National Park Service and Bureau of Land Management for granting us permission to study the cyanobacterial flora within Joshua Tree National Park (permit JOTR-2006-SCI-0018), Mojave Desert National Preserve (MOJA-2008-SCI-0024 and MOJA-2009-SCI-0039), and Grand Staircase-Escalante National Monument (permit UT-06-032-12-P). National Science Foundation (NSF) funded J.R. Johansen’s sample collection of soils from the Atacama Desert, isolation, and sequencing of those strains (NSF grant numbers DEB-0842702 and DEB-841734, respectively), as well as the collection and biodiversity research of biological soil crusts from various North American desert locations (NSF grant number DEB-9870201). This work is supported by the USDA National Institute of Food and Agriculture and Hatch Appropriations under Project #PEN04949 to E.C and Hatch project CA-R-PPA-5062-H to J.E.S. Any opinions, findings, conclusions, or recommendations expressed in this material are those of the authors and do not necessarily reflect the views of the National Science Foundation. The California Desert Research Fund at The Community Foundation, Robert Lee Graduate Student Research Grant, and the Phycological Society Grants in Aid of Research fund awarded to Nicole Pietrasiak provided support for the sampling campaigns and subsequent cyanobacterial research associated with strains from Joshua Tree National Park and Mojave Desert National Preserve. J.E.S. is a CIFAR Fellow in the program Fungal Kingdom: Threats and Opportunities. John Carroll University supported J.R.J.’s cyanobacterial culture collection since 1996. We greatly acknowledge the numerous students and colleagues who over the past 30 years collaborated with Johansen and led to the isolation of many of the strains investigated in this study.

## Supplementary

**Table S1:** Metadata for the cyanobacteria used in this study. The “Map ID” column corresponds with the locations of the cyanobacteria depicted in Figure 1. The annual mean temperature and annual precipitation were derived from BioClim data when accurate latitude and longitude coordinates were provided for the cyanobacterial hosts. The * denote cyanobacteria that were not previously published in Ward et al. 2021.

**Table S2:** Summary statistics from each MAG including completeness, contamination, number of contigs, genome size, and taxonomic information.

## References

1. Büdel B. 2011. Cyanobacteria: Habitats and Species, p. 11–21. In Lüttge, U, Beck, E, Bartels, D (eds.), Plant Desiccation Tolerance. Springer, Berlin, Heidelberg.

2. Ward RD, Stajich JE, Johansen JR, Huntemann M, Clum A, Foster B, Foster B, Roux S, Palaniappan K, Varghese N, Mukherjee S, Reddy TBK, Daum C, Copeland A, Chen I-MA, Ivanova NN, Kyrpides NC, Shapiro N, Eloe-Fadrosh EA, Pietrasiak N. 2021. Metagenome Sequencing to Explore Phylogenomics of Terrestrial Cyanobacteria. Microbiology Resource Announcements 10:10.1128/mra.00258-21.

3. Steiner JL, Brandani CB, Chappell A, Castaño-Sanchez J, Hoellrich M, McIntosh MM, Nyamuryekung’e S, Pietrasiak N, Rotz A, Webb NP. 2023. Distinctive Dryland Soil Carbon Transformations: Insights from Arid Rangelands of SW United StatesSoil and Drought. CRC Press.

4. Pankratova EM. 2006. Functioning of cyanobacteria in soil ecosystems. Eurasian Soil Sc 39:S118–S127.

5. Sepehr A, Hassanzadeh M, Rodriguez-Caballero E. 2019. The protective role of cyanobacteria on soil stability in two Aridisols in northeastern Iran. Geoderma Regional 16:e00201.

6. Rodriguez-Caballero E, Belnap J, Büdel B, Crutzen PJ, Andreae MO, Pöschl U, Weber B. 2018. Dryland photoautotrophic soil surface communities endangered by global change. Nature Geoscience 11:185–189.

7. Kneitel JM, Lessin CL. 2010. Ecosystem-phase interactions: aquatic eutrophication decreases terrestrial plant diversity in California vernal pools. Oecologia 163:461–469.

8. Vigil JP, Schuler MS. 2024. Salt pollution reduces turbidity, dissolved organic matter, and cyanobacteria in experimental vernal pool communities. Science of The Total Environment 931:172948.

9. Pentecost A, Whitton BA. 2012. Subaerial Cyanobacteria, p. 291–316. In Whitton, BA (ed.), Ecology of Cyanobacteria II: Their Diversity in Space and Time. Springer Netherlands, Dordrecht.

10. Hauer T, Mühlsteinová R, Bohunická M, Kaštovský J, Mareš J. 2015. Diversity of cyanobacteria on rock surfaces. Biodivers Conserv 24:759–779.

11. Villa F, Cappitelli F. 2019. The Ecology of Subaerial Biofilms in Dry and Inhospitable Terrestrial Environments. 10. Microorganisms 7:380.

12. Ortega-Calvo JJ, Ariño X, Hernandez-Marine M, Saiz-Jimenez C. 1995. Factors affecting the weathering and colonization of monuments by phototrophic microorganisms. Science of The Total Environment 167:329–341.

13. Couradeau E, Giraldo-Silva A, De Martini F, Garcia-Pichel F. 2019. Spatial segregation of the biological soil crust microbiome around its foundational cyanobacterium, Microcoleus vaginatus, and the formation of a nitrogen-fixing cyanosphere. Microbiome 7:55.

14. Nelson C, Giraldo-Silva A, Garcia-Pichel F. 2021. A symbiotic nutrient exchange within the cyanosphere microbiome of the biocrust cyanobacterium, Microcoleus vaginatus. The ISME Journal 15:282–292.

14. Barea J-M, Pozo M-J, Azcón R, Azcón-Aguilar C. 2013. Microbial Interactions in the Rhizosphere, p. 29–44. In Molecular Microbial Ecology of the Rhizosphere. John Wiley & Sons, Ltd.

15. Zheng Q, Hu Y, Kosina SM, Van Goethem MW, Tringe SG, Bowen BP, Northen TR. 2023. Conservation of beneficial microbes between the rhizosphere and the cyanosphere. New Phytologist 240:1246–1258.

16. Nelson C, Garcia-Pichel F. 2021. Beneficial Cyanosphere Heterotrophs Accelerate Establishment of Cyanobacterial Biocrust. Applied and Environmental Microbiology 87:e01236–21.

17. Nelson C, Dadi P, Shah DD, Garcia-Pichel F. 2024. Spatial organization of a soil cyanobacterium and its cyanosphere through GABA/Glu signaling to optimize mutualistic nitrogen fixation. The ISME Journal wrad029.

18. Pascault N, Rué O, Loux V, Pédron J, Martin V, Tambosco J, Bernard C, Humbert J, Leloup J. 2021. Insights into the cyanosphere: capturing the respective metabolisms of cyanobacteria and chemotrophic bacteria in natural conditions? Environmental Microbiology Reports 13:364–374.

19. Carmichael W. 1986. Isolation, culture, and toxicity testing of toxic freshwater cyanobacteria (blue-green algae). Fundamental Research in Homogenous Catalysis 3:1249–1262.

20. Arkin AP, Cottingham RW, Henry CS, Harris NL, Stevens RL, Maslov S, Dehal P, Ware D, Perez F, Canon S, Sneddon MW, Henderson ML, Riehl WJ, Murphy-Olson D, Chan SY, Kamimura RT, Kumari S, Drake MM, Brettin TS, Glass EM, Chivian D, Gunter D, Weston DJ, Allen BH, Baumohl J, Best AA, Bowen B, Brenner SE, Bun CC, Chandonia J-M, Chia J-M, Colasanti R, Conrad N, Davis JJ, Davison BH, DeJongh M, Devoid S, Dietrich E, Dubchak I, Edirisinghe JN, Fang G, Faria JP, Frybarger PM, Gerlach W, Gerstein M, Greiner A, Gurtowski J, Haun HL, He F, Jain R, Joachimiak MP, Keegan KP, Kondo S, Kumar V, Land ML, Meyer F, Mills M, Novichkov PS, Oh T, Olsen GJ, Olson R, Parrello B, Pasternak S, Pearson E, Poon SS, Price GA, Ramakrishnan S, Ranjan P, Ronald PC, Schatz MC, Seaver SMD, Shukla M, Sutormin RA, Syed MH, Thomason J, Tintle NL, Wang D, Xia F, Yoo H, Yoo S, Yu D. 2018. KBase: The United States Department of Energy Systems Biology Knowledgebase. Nat Biotechnol 36:566–569.

21. Andrews S. 2017. FastQC: a quality control tool for high throughput sequence data. 2010

22. Bolger AM, Lohse M, Usadel B. 2014. Trimmomatic: a flexible trimmer for Illumina sequence data. Bioinformatics 30:2114–2120.

23. Nurk S, Meleshko D, Korobeynikov A, Pevzner PA. 2017. metaSPAdes: a new versatile metagenomic assembler. Genome Res 27:824–834.

24. Wu Y-W, Simmons BA, Singer SW. 2016. MaxBin 2.0: an automated binning algorithm to recover genomes from multiple metagenomic datasets. Bioinformatics 32:605–607.

25. Chaumeil P-A, Mussig A, Philip H, Parks D. 2019. GTDB-Tk: a toolkit to classify genomes with the Genome Taxonomy Database. Bioinformatics (Oxford, England) 36.

26. Bowers RM, Kyrpides NC, Stepanauskas R, Harmon-Smith M, Doud D, Reddy TBK, Schulz F, Jarett J, Rivers AR, Eloe-Fadrosh EA, Tringe SG, Ivanova NN, Copeland A, Clum A, Becraft ED, Malmstrom RR, Birren B, Podar M, Bork P, Weinstock GM, Garrity GM, Dodsworth JA, Yooseph S, Sutton G, Glöckner FO, Gilbert JA, Nelson WC, Hallam SJ, Jungbluth SP, Ettema TJG, Tighe S, Konstantinidis KT, Liu W-T, Baker BJ, Rattei T, Eisen JA, Hedlund B, McMahon KD, Fierer N, Knight R, Finn R, Cochrane G, Karsch-Mizrachi I, Tyson GW, Rinke C, Lapidus A, Meyer F, Yilmaz P, Parks DH, Murat Eren A, Schriml L, Banfield JF, Hugenholtz P, Woyke T. 2017. Minimum information about a single amplified genome (MISAG) and a metagenome-assembled genome (MIMAG) of bacteria and archaea. Nat Biotechnol 35:725–731.

27. Andersen KS, Kirkegaard RH, Karst SM, Albertsen M. 2018. ampvis2: An R package to analyse and visualise 16S rRNA amplicon data. BioRxiv 299537.

28. Liu C, Cui Y, Li X, Yao M. 2021. microeco: An R package for data mining in microbial community ecology. FEMS Microbiology Ecology 97:fiaa255.

29. Karger DN, Conrad O, Böhner J, Kawohl T, Kreft H, Soria-Auza RW, Zimmermann NE, Linder HP, Kessler M. 2017. Climatologies at high resolution for the earth’s land surface areas. Sci Data 4:170122.

30. Kolde R, Kolde MR. 2015. Package ‘pheatmap.’ R package 1:790.

31. D Ainsworth T, Krause L, Bridge T, Torda G, Raina J-B, Zakrzewski M, Gates RD, Padilla-Gamiño JL, Spalding HL, Smith C, Woolsey ES, Bourne DG, Bongaerts P, Hoegh-Guldberg O, Leggat W. 2015. The coral core microbiome identifies rare bacterial taxa as ubiquitous endosymbionts. The ISME Journal 9:2261–2274.

32. Sweet MJ, Brown BE, Dunne RP, Singleton I, Bulling M. 2017. Evidence for rapid, tide-related shifts in the microbiome of the coral Coelastrea aspera. Coral Reefs 36:815–828.

33. Pham VHT, Jeong S, Chung S, Kim J. 2016. Brevundimonas albigilva sp. nov., isolated from forest soil. International Journal of Systematic and Evolutionary Microbiology 66:1144–1150.

34. Dahal RH, Kim J. 2018. Brevundimonas humi sp. nov., an alphaproteobacterium isolated from forest soil. International Journal of Systematic and Evolutionary Microbiology 68:709– 714.

35. Qu J-H, Fu Y-H, Li X-D, Li H-F, Tian H-L. 2019. Brevundimonas lutea sp. nov., isolated from lake sediment. International Journal of Systematic and Evolutionary Microbiology 69:1417–1422.

36. Naqqash T, Imran A, Hameed S, Shahid M, Majeed A, Iqbal J, Hanif MK, Ejaz S, Malik KA. 2020. First report of diazotrophic Brevundimonas spp. as growth enhancer and root colonizer of potato. Sci Rep 10:12893.

37. Ryu SH, Park M, Lee JR, Yun P-Y, Jeon CO. 2007. Brevundimonas aveniformis sp. nov., a stalked species isolated from activated sludge. International Journal of Systematic and Evolutionary Microbiology 57:1561–1565.

38. Yoon J-H, Kang S-J, Oh HW, Lee J-S, Oh T-K. 2006. Brevundimonas kwangchunensis sp. nov., isolated from an alkaline soil in Korea. International Journal of Systematic and Evolutionary Microbiology 56:613–617.

39. Tóth E, Szuróczki S, Kéki Zs, Kosztik J, Makk J, Bóka K, Spröer C, Márialigeti K, Schumann P. 2017. Brevundimonas balnearis sp. nov., isolated from the well water of a thermal bath. International Journal of Systematic and Evolutionary Microbiology 67:1033– 1038.

40. Yoo S-H, Weon H-Y, Kim B-Y, Hong S-B, Kwon S-W, Cho Y-H, Go S-J, Stackebrandt E. 2006. Devosia soli sp. nov., isolated from greenhouse soil in Korea. International Journal of Systematic and Evolutionary Microbiology 56:2689–2692.

41. Yoon J-H, Kang S-J, Park S, Oh T-K. 2007. Devosia insulae sp. nov., isolated from soil, and emended description of the genus Devosia. International Journal of Systematic and Evolutionary Microbiology 57:1310–1314.

42. Jia Y-Y, Sun C, Pan J, Zhang W-Y, Zhang X-Q, Huo Y-Y, Zhu X-F, Wu M. 2014. Devosia pacifica sp. nov., isolated from deep-sea sediment. International Journal of Systematic and Evolutionary Microbiology 64:2637–2641.

43. Chen Y, Zhu S, Lin D, Wang X, Yang J, Chen J. 2019. Devosia naphthalenivorans sp. nov., isolated from East Pacific Ocean sediment. International Journal of Systematic and Evolutionary Microbiology 69:1974–1979.

44. Ma F, Zi Z-D, Li W, Wang Z-X, Lu J, Lv J. 2021. Devosia sediminis sp. nov., isolated from subterranean sediment. Arch Microbiol 203:4517–4523.

45. Rivas R, Velázquez E, Willems A, Vizcaíno N, Subba-Rao NS, Mateos PF, Gillis M, Dazzo FB, Martínez-Molina E. 2002. A New Species of Devosia That Forms a Unique Nitrogen-Fixing Root-Nodule Symbiosis with the Aquatic Legume Neptunia natans (L.f.) Druce. Applied and Environmental Microbiology 68:5217–5222.

46. Kim JM, Baek W, Choi BJ, Bayburt H, Baek JH, Han DM, Lee SC, Jeon CO. 2024. Devosia rhodophyticola sp. nov. and Devosia algicola sp. nov., isolated from a marine red alga. International Journal of Systematic and Evolutionary Microbiology 74:006223.

47. Talwar C, Nagar S, Kumar R, Scaria J, Lal R, Negi RK. 2020. Defining the Environmental Adaptations of Genus Devosia: Insights into its Expansive Short Peptide Transport System and Positively Selected Genes. Sci Rep 10:1151.

48. Godoy F, Vancanneyt M, Martínez M, Steinbüchel A, Swings J, Rehm BHA. 2003. Sphingopyxischilensis sp. nov., a chlorophenol-degrading bacterium that accumulates polyhydroxyalkanoate, and transfer of Sphingomonas alaskensis to Sphingopyxis alaskensis comb. nov. International Journal of Systematic and Evolutionary Microbiology 53:473–477.

49. Zhang D-C, Liu H-C, Xin Y-H, Zhou Y-G, Schinner F, Margesin R. 2010. Sphingopyxis bauzanensis sp. nov., a psychrophilic bacterium isolated from soil. International Journal of Systematic and Evolutionary Microbiology 60:2618–2622.

50. Kaminski MA, Sobczak A, Spolnik G, Danikiewicz W, Dziembowski A, Lipinski L. 2018. Sphingopyxis lindanitolerans sp. nov. strain WS5A3pT enriched from a pesticide disposal site. International Journal of Systematic and Evolutionary Microbiology 68:3935–3941.

51. Takeuchi M, Hamana K, Hiraishi A. 2001. Proposal of the genus Sphingomonas sensu stricto and three new genera, Sphingobium, Novosphingobium and Sphingopyxis, on the basis of phylogenetic and chemotaxonomic analyses. International Journal of Systematic and Evolutionary Microbiology 51:1405–1417.

52. Sharma M, Khurana H, Singh DN, Negi RK. 2021. The genus *Sphingopyxis*: Systematics, ecology, and bioremediation potential - A review. Journal of Environmental Management 280:111744.

53. Lindström K, Mousavi SA. 2020. Effectiveness of nitrogen fixation in rhizobia. Microbial Biotechnology 13:1314–1335.

54. Thomas L, Rahman Z. 2020. A Genome-Wide Investigation on Symbiotic Nitrogen-Fixing Bacteria in Leguminous Plants, p. 55–73. In Varma, A, Tripathi, S, Prasad, R (eds.), Plant Microbe Symbiosis. Springer International Publishing, Cham.

55. Carvalho FM, Souza RC, Barcellos FG, Hungria M, Vasconcelos ATR. 2010. Genomic and evolutionary comparisons of diazotrophic and pathogenic bacteria of the order Rhizobiales. BMC Microbiol 10:37.

56. Ling N, Wang T, Kuzyakov Y. 2022. Rhizosphere bacteriome structure and functions. Nat Commun 13:836.

57. Vieira S, Sikorski J, Dietz S, Herz K, Schrumpf M, Bruelheide H, Scheel D, Friedrich MW, Overmann J. 2020. Drivers of the composition of active rhizosphere bacterial communities in temperate grasslands. The ISME Journal 14:463–475.

58. Sweeney CJ, de Vries FT, van Dongen BE, Bardgett RD. 2021. Root traits explain rhizosphere fungal community composition among temperate grassland plant species. New Phytologist 229:1492–1507.

59. Vaccarino MA, Johansen JR. 2012. Brasilonema Angustatum Sp. Nov. (nostocales), a New Filamentous Cyanobacterial Species from the Hawaiian Islands. Journal of Phycology 48:1178–1186.

60. Renaudin M, Laforest-Lapointe I, Bellenger J-P. 2022. Unraveling global and diazotrophic bacteriomes of boreal forest floor feather mosses and their environmental drivers at the ecosystem and at the plant scale in North America. Science of The Total Environment 837:155761.

61. Hungria M, Menna P, Delamuta JRM. 2015. Bradyrhizobium, the Ancestor of All Rhizobia: Phylogeny of Housekeeping and Nitrogen-Fixation Genes, p. 191–202. In Biological Nitrogen Fixation. John Wiley & Sons, Ltd.

62. Kuykendall LD. 2015. Bradyrhizobium. Bergey’s Manual of Systematics of Archaea and Bacteria 1–11.

63. Zhu C, Friman V-P, Li L, Xu Q, Guo J, Guo S, Shen Q, Ling N. 2022. Meta-analysis of diazotrophic signatures across terrestrial ecosystems at the continental scale. Environmental Microbiology 24:2013–2028.

64. Sutherland IW. 1995. Polysaccharide lyases. FEMS Microbiology Reviews 16:323–347.

65. Kennedy L, Sutherland IW. 1996. Polysaccharide lyases from gellan-producing Sphingomonas spp. Microbiology 142:867–872.

66. Hashimoto W, Miyake O, Ochiai A, Murata K. 2005. Molecular identification of Sphingomonas sp. A1 Alginate lyase (A1-IV′) as a member of novel polysaccharide lyase family 15 and implications in alginate lyase evolution. Journal of Bioscience and Bioengineering 99:48–54.

67. Guo Q, Dan M, Zheng Y, Shen J, Zhao G, Wang D. 2023. Improving the thermostability of a novel PL-6 family alginate lyase by rational design engineering for industrial preparation of alginate oligosaccharides. International Journal of Biological Macromolecules 249:125998.

68. Khan F, Akhlaq A, Rasool MH, Srinuanpan S. 2024. Cyanobacterial Bioactive Compounds: Synthesis, Extraction, and Applications, p. 215–243. In Mehmood, MA, Verma, P, Shah, MP, Betenbaugh, MJ (eds.), Pharmaceutical and Nutraceutical Potential of Cyanobacteria. Springer International Publishing, Cham.

69. Warshan D, Espinoza JL, Stuart RK, Richter RA, Kim S-Y, Shapiro N, Woyke T, C Kyrpides N, Barry K, Singan V, Lindquist E, Ansong C, Purvine SO, M Brewer H, Weyman PD, Dupont CL, Rasmussen U. 2017. Feathermoss and epiphytic Nostoc cooperate differently: expanding the spectrum of plant–cyanobacteria symbiosis. The ISME Journal 11:2821–2833.

70. Payasi A, Sanwal R, Sanwal GG. 2009. Microbial pectate lyases: characterization and enzymological properties. World J Microbiol Biotechnol 25:1–14.

71. Xie F, Murray JD, Kim J, Heckmann AB, Edwards A, Oldroyd GED, Downie JA. 2012. Legume pectate lyase required for root infection by rhizobia. Proceedings of the National Academy of Sciences 109:633–638.

72. Lan S, Zhang Q, Wu L, Liu Y, Zhang D, Hu C. 2014. Artificially Accelerating the Reversal of Desertification: Cyanobacterial Inoculation Facilitates the Succession of Vegetation Communities. Environ Sci Technol 48:307–315.

73. Chamizo S, Mugnai G, Rossi F, Certini G, De Philippis R. 2018. Cyanobacteria inoculation improves soil stability and fertility on different textured soils: gaining insights for applicability in soil restoration. Frontiers in Environmental Science 6:49.

74. Muñoz-Rojas M, Chilton A, Liyanage GS, Erickson TE, Merritt DJ, Neilan BA, Ooi MKJ. 2018. Effects of indigenous soil cyanobacteria on seed germination and seedling growth of arid species used in restoration. Plant and Soil 429:91–100.

75. Yadav P, Singh RP, Rana S, Joshi D, Kumar D, Bhardwaj N, Gupta RK, Kumar A. 2022. Mechanisms of Stress Tolerance in Cyanobacteria under Extreme Conditions. 4. Stresses 2:531–549.

76. Eldridge DJ, Reed S, Travers SK, Bowker MA, Maestre FT, Ding J, Havrilla C, Rodriguez-Caballero E, Barger N, Weber B, Antoninka A, Belnap J, Chaudhary B, Faist A, Ferrenberg S, Huber-Sannwald E, Malam Issa O, Zhao Y. 2020. The pervasive and multifaceted influence of biocrusts on water in the world’s drylands. Global Change Biology 26:6003– 6014.

77. Jafarpoor A, Sadeghi SH, Darki BZ, Homaee M. 2022. Changes in morphologic, hydraulic, and hydrodynamic properties of rill erosion due to surface inoculation of endemic soil cyanobacteria. Catena 208:105782.

